# Correlation between structure and function in phosphatidylinositol lipid-dependent Kir2.2 gating

**DOI:** 10.1101/2021.02.15.431350

**Authors:** Yuxi Zhang, Xiao Tao, Roderick MacKinnon

## Abstract

Inward rectifier K^+^(Kir) channels regulate cell membrane potential. Different Kir channels respond to unique ligands, but all are regulated by phosphatidylinositol 4,5-bisphosphate (PI(4,5)P_2_). Using planar lipid bilayers we show that Kir2.2 exhibits bursts of openings separated by long quiescent inter-burst periods. Increasing PI(4,5)P_2_ concentration shortens the Kir2.2 inter-burst duration and lengthens the burst duration without affecting dwell times within a burst. From this, we propose that burst and inter-burst durations correspond to the CTD-docked and CTD-undocked conformations observed in the presence and absence of PI(4,5)P_2_ in atomic structures. We also studied the effect of different phosphatidylinositol lipids on Kir2.2 activation and conclude that the 5’ phosphate is essential to Kir2.2 pore opening. Other phosphatidylinositol lipids can compete with PI(4,5)P_2_ but cannot activate Kir2.2 without the 5’ phosphate. PI(4)P, which is directly interconvertible to and from PI(4,5)P_2_, might thus be a regulator of Kir channels in the plasma membrane.

## Introduction

Inward rectifier K^+^ (Kir) channels are so named because they conduct K^+^ better at negative membrane potentials, allowing them to affect ‘resting’ potential while minimizing K^+^ outflow during depolarization (Hagiwara et al., 1976; Hagiwara and Takahashi, 1974; Hille, 2001; Hodgkin AL, 1949; Hodgkin and Horowicz, 1959; Katz, 1949). Kir channels are involved in many physiological processes, including the regulation of cell membrane potential, cellular pacemaker activity and hormone secretion (Hibino et al., 2010; Hille, 2001). Structurally, Kir channels comprise four subunits, each containing a transmembrane domain (TMD) with a selectivity filter and a cytoplasmic domain (CTD) connected to the TMD by a linker (Tao et al., 2009; Whorton and MacKinnon, 2011). While different subclasses of Kir channels are regulated by unique modulators, for example G proteins in the GIRK channel and ATP in the KATP channel, all Kir channels are regulated by PI(4,5)P_2_, a signaling lipid present in the cell plasma membrane (Ashcroft, 1988; Hibino et al., 2010; Hilgemann et al., 2001; Huang et al., 1998; Logothetis et al., 1987; Nichols and Lederer, 1991; Stanfield et al., 2002).

X-ray crystallographic and cryo-electron microscopic studies have shown that PI(4,5)P_2_ can modify the conformation of Kir channels (Hansen et al., 2011; Niu et al., 2020; Tao et al., 2009; Whorton and MacKinnon, 2011). For example, in Kir2.2 in the absence of PI(4,5)P_2_ the CTD disengages from the TMD to form a “CTD-undocked” conformation, which is accompanied by a tightly constricted inner helix gate (Tao et al., 2009). Upon PI(4,5)P_2_ binding, the CTD engages the TMD to form a “CTD-docked” conformation and the inner helix gate widens (Figure 1A) (Hansen et al., 2011). A similar PI(4,5)P_2_-mediated conformational change was observed in GIRK, and structures determined under varying PI(4,5)P_2_ concentrations indicate that PI(4,5)P_2_ concentrations regulate the equilibrium distribution among CTD-docked and CTD-undocked conformations (Niu et al., 2020).

**Figure 1.**
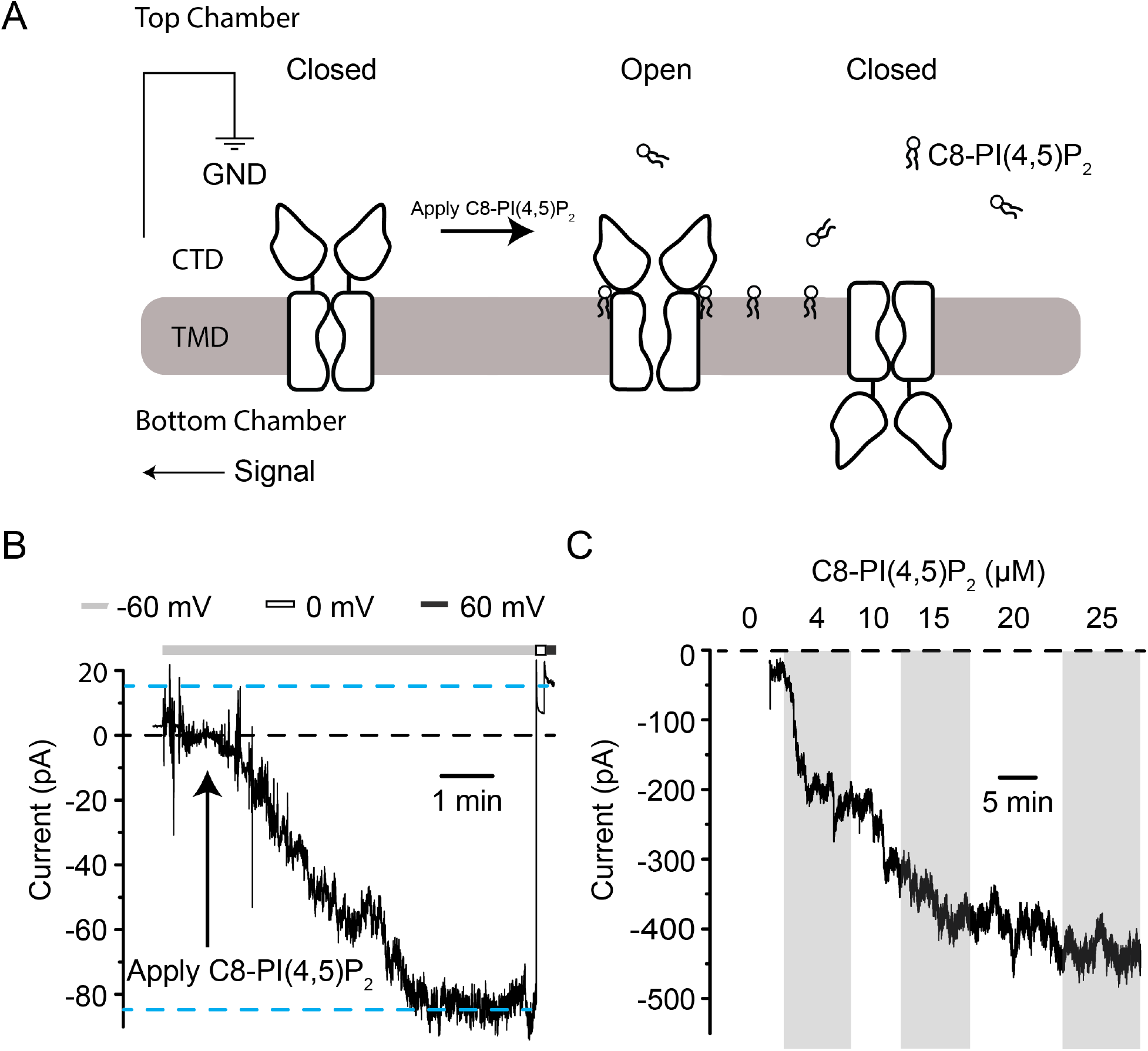
PIP_2_-dependent activation of Kir2.2 reconstituted into planar lipid bilayer. (A) Schematic of the planar lipid bilayer system. (B) C8-PI(4,5)P_2_ is both necessary and sufficient for Kir2.2 channel activation. Representative current trace recorded at indicated voltages before and after addition of 10 μM C8-PI(4,5)P_2_ to the top chamber is shown. Zero-current level is indicated by black dashed line. Amplitude of the current at −60 mV and 60 mV (cyan dashed lines) shows strong rectification. The current was inverted to follow electrophysiological convention. (C) Current of Kir2.2 in responding to increasing C8-PI(4,5)P_2_ concentrations. Increasing the amount of C8-PI(4,5)P_2_ (concentration indicated above) was applied to the top chamber followed by mixing. The membrane was held at −100 mV.

Previous electrophysiological studies using inside-out patches from cell membranes showed that two different Kir channels, the KATP channel and a G protein-independent mutant of the GIRK channel, both gate in bursts, that is, intervals of rapid channel opening and closing were separated by long quiescent periods (Enkvetchakul et al., 2000; Jin et al., 2008). Furthermore, in the GIRK channel, PI(4,5)P_2_ influenced the duration of the burst periods (Jin et al., 2008). In the KATP channel, PI(4,5)P_2_ influenced the duration of quiescent periods, without changing the kinetics within the burst (Enkvetchakul et al., 2000). Two other Kir channels, Kir2.1 and Kir2.2, were studied following purification and reconstitution in lipid vesicles (D’Avanzo et al., 2010). Using Rb^+^ flux and patch clamp analysis these channels were found to be activated by PI(4,5)P_2_ and other phosphatidylinositol lipid derivatives in the absence of other regulatory molecules.

In this paper we present an analysis of the Kir2.2 channel gating response to phosphatidylinositol lipids using the planar lipid bilayer recording system. This system offers complete chemical control of lipid, solution and protein composition as well as free access to the solution bathing the surfaces of the membrane (Miller and Racker, 1976; Montal and Mueller, 1972; Wang et al., 2014). Given that we already have a detailed description of the structural changes that Kir2.2 undergoes upon binding of PI(4,5)P_2_ (Hansen et al., 2011; Tao et al., 2009), our goal here is to correlate PI(4,5)P_2_-dependent gating properties with the known structural changes. Furthermore, given our detailed chemical knowledge of the PI(4,5)P_2_ binding site on Kir2.2, we characterize and interpret the influence of different phosphatidylinositol lipid derivatives on channel gating. We find that competition for the PI(4,5)P_2_ binding site by PI(4)P inhibits Kir2.2 activity. Because both PI(4)P and PI(4,5)P_2_ contribute substantially to the pool of plasma membrane phosphatidylinositol lipids, their competition and interconversion might be relevant to Kir2.2 *in vivo* (Di Paolo and De Camilli, 2006; Hammond and Balla, 2015; Logothetis et al., 2015).

## Results

### Dependence of Kir2.2 opening on C8-PI(4,5)P_2_ concentration

Figure 1A shows a schematic of the planar lipid bilayer system used in this study (Miller and Racker, 1976; Montal and Mueller, 1972; Wang et al., 2014). A lipid bilayer with a defined composition separates the top and bottom chambers, each filled with electrolyte solution. Liposomes containing Kir2.2 channels are fused with the bilayer, resulting in some channels with the CTD facing the top chamber and others with the CTD facing the bottom chamber (Figure 1A). Soluble reagents such as C8-PI(4,5)P_2_ (PI(4,5)P_2_ with 8-carbon acyl chains) are added to the top chamber. Because C8-PI(4,5)P_2_ is membrane-impermeant, only channels with their CTD facing the top chamber are activated (Figure 1A) (Wang et al., 2014).

We first looked at the dependence of channel activity on PI(4,5)P_2_ concentration. In the absence of C8-PI(4,5)P_2_, no channel openings were observed. Upon addition of C8-PI(4,5)P_2_, current exhibiting strong inward rectification was induced (Figure 1B), indicating that PI(4,5)P_2_ is both necessary and sufficient for channel opening, as was found previously in a different reconstitution system (D’Avanzo et al., 2010). Kir2.2 activity rises steeply over the C8-PI(4,5)P_2_ concentration range 0 to 10 μM and approaches maximal activation by 15 μM (Figure 1C). Notably, C8-PI(4,5)P_2_-induced current begins to decrease spontaneously after about 30 min (Figure 1 – figure supplement 1). The disappearance of channel activity over time introduces uncertainty to the concentration-dependence of Kir2.2 activity and imposes a limitation on the kinetic analysis, described below.

### Influence of C8-PI(4,5)P_2_ on the gating kinetics of Kir2.2

We next looked at how C8-PI(4,5)P_2_ influences the kinetics of Kir2.2 gating. Figure 2A shows a single-channel trace recorded in the presence of 3, 6 and 15 μM C8-PI(4,5)P_2_. The channel opens in bursts of activity separated by quiescent intervals. We see from this trace two processes that operate on very different timescales. The relatively fast process, occurring on the sub-second timescale, accounts for rapid channel opening and closing within a burst of activity (Figure 2A, inset). The slow process, occurring on the minute timescale, accounts for the appearance and disappearance of bursts. Lifetime histograms for events within bursts show single open-time and single closed-time distributions, which correspond to the relatively rapid gating transitions that occur within a burst. (Figure 2 A-E). Note that the histograms are essentially unchanged when the concentration of C8-PI(4,5)P_2_ is increased from 6 μM to 15 μM (compare Figure 2 B with D and Figure 2 C with E). This observation indicates that gating transitions within a burst are insensitive to the C8-PI(4,5)P_2_ concentration over a range that influences the open probability (Figure 1C). A second, longer closed dwell time exists because we see it in the raw trace (Figure 2A), however, its frequency is too low to accumulate enough events during the recording, which is limited in duration owing to the phenomenon of channel disappearance over time. Figure 2F shows the connectivity diagram for two closed and one open state (Figure 2F, 1). One linear kinetic sub-scheme, 2, is incompatible with the channel record because openings would not be interrupted by brief closures. The remaining two linear kinetic sub-schemes, 3 and 4, are compatible with the record, which cannot distinguish among them.

**Figure 2.**
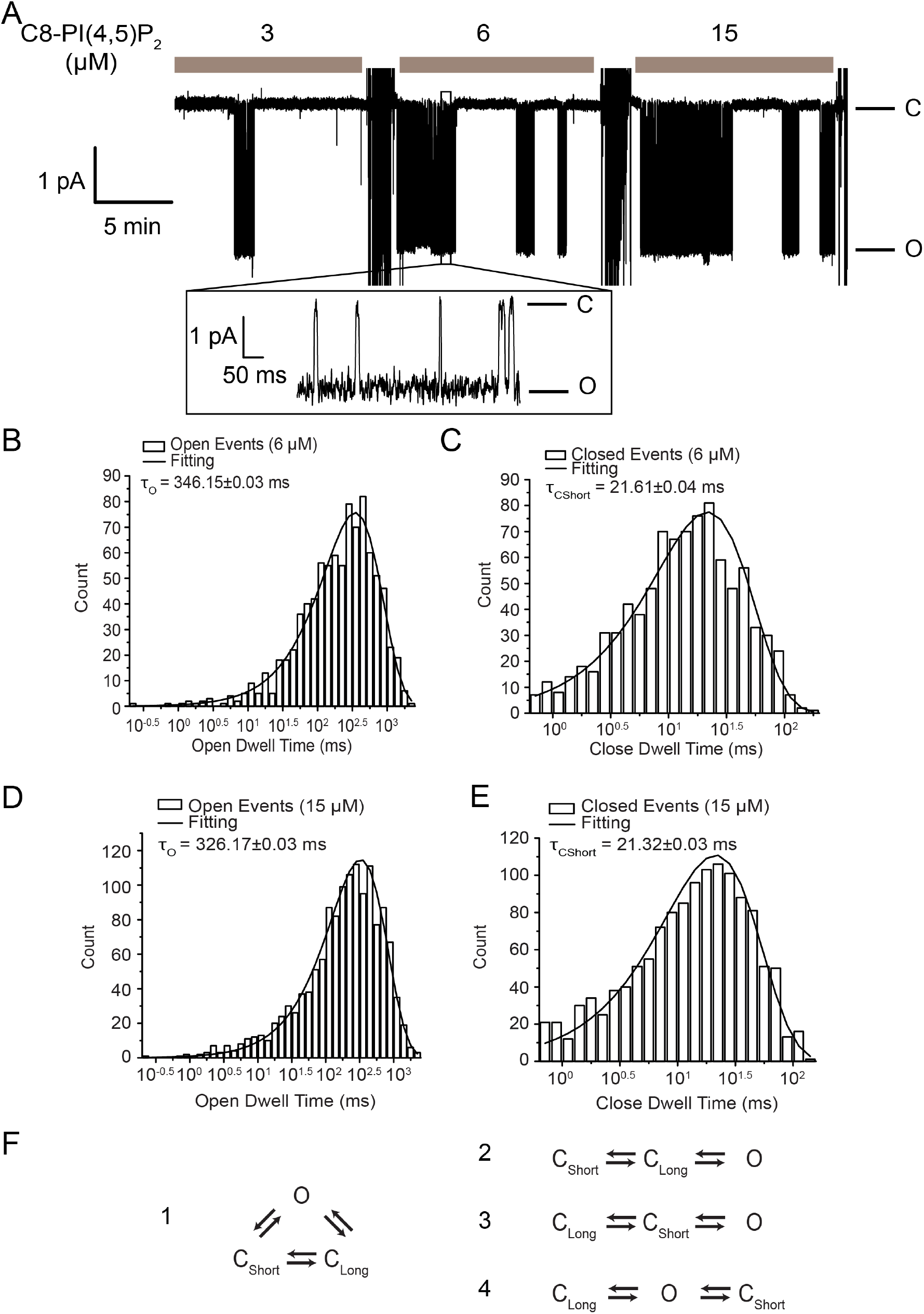
Single-channel analysis of Kir2.2. (A) Single-channel recording of Kir2.2 at −100 mV in the presence of 3, 6 and 15 μM C8-PI(4,5)P_2_. An expanded trace is shown in inset. The open (o) and closed (c) levels are indicated. Large current deflections between adjacent C8-PI(4,5)P_2_ concentrations are due to mixing of the solution after each addition of C8-PI(4,5)P_2_. (B) The open dwell time distribution of Kir2.2 at 6 μM C8-PI(4,5)P_2_ from panel (A) is plotted and fit with a single exponential (solid line). (C) The dwell time distribution of closed events within the burst at 6 μM C8-PI(4,5)P_2_ from panel (A) is plotted and fit with a single exponential (solid line). (D) The open dwell time distribution of Kir2.2 at 15 μM C8-PI(4,5)P_2_ from panel (A) is plotted and fit with a single exponential (solid line). (E) The dwell time distribution of closed events within the burst at 15 μM C8-PI(4,5)P_2_ from panel (A) is plotted and fit with a single exponential (solid line). (F) Connectivity diagram (1) and possible linear kinetic sub-schemes (2-4) with one short closed state (C_Short_), one long closed state (C_Long_) and one open state (O).

In the context of the compatible kinetic schemes, we next ask which transitions are affected by the concentration of C8-PI(4,5)P_2_? Given the low frequency of inter-burst intervals, confounded by channel disappearance over time, we resorted to studying bilayer membranes with several channels present at once. While this approach is not ideal, it provides a sufficient number of events to estimate the rate constants. Figure 3A shows a multi-channel membrane in the presence of 3, 6 and 15 μM C8-PI(4,5)P_2_. The 15 μM record was used to estimate the total number of channels in the membrane (see Methods), while 3 and 6 μM records were subject to kinetic analysis (Table 1) (Csanady, 2000). The analysis assumes that all channels are identical in their behavior. For most channel types we have studied, including Kir2.2, this assumption is only approximately true. There appeared to be a small fraction of outlier channels with lower or higher than average open probability, which undoubtedly contributed to the variation in rate constant values between different experiments. This limitation notwithstanding, Table 1 shows that C8-PI(4,5)P_2_ affects only the rate constants for transitions into and out of C_Long_. In detail, when C8-PI(4,5)P_2_ is increased, the burst periods lengthen and the quiescent periods shorten. Consistent with the single-channel trace in Figure 2, rate constants for opening and closing within a burst are insensitive to C8-PI(4,5)P_2_ concentration: within a burst, only the mean number of transitions is affected by C8-PI(4,5)P_2_. Table 1 reports rate constant values for scheme 3, but scheme 4 (Figure 2F) would yield a similar conclusion, that only rate constants into and out of C_Long_ are sensitive to C8-PI(4,5)P_2_ concentration.

**Table 1.** Kinetics of Kir2.2 in the presence of 3 and 6 μM C8-PI(4,5)P_2_. Kinetics of Kir2.2 based on scheme 3 (C_Long_ == C_Short_ == O) in the presence of 3 and 6 μM C8-PI(4,5)P_2_. Rate constant (k), open probability (P(o)), mean burst duration (T_burst_), mean inter-burst interval (T_inter-burst_), mean duration of C_Short_ state and mean number of short closures per burst (#C_Short_ per burst) are calculated from 3 different recordings.

| Recording Number | Channel Number | C8-PI(4,5)$\text{P}_2$($\mu\text{M}$) | $k_{12}(\text{s}^{-1})$ | $k_{21}(\text{s}^{-1})$ | $k_{23}(\text{s}^{-1})$ | $k_{32}(\text{s}^{-1})$ | P(o) | $T_{\text{burst}}(\text{s})$ | $T_{\text{inter-burst}}(\text{s})$ | Mean Duration of $\text{C}_{\text{Short}}$ (ms) | #$\text{C}_{\text{Short}}$ per burst |
| --- | --- | --- | --- | --- | --- | --- | --- | --- | --- | --- | --- |
| 1 | 8 | 3 | 0.0097<br>$\pm 0.0007$ | 1.7$\pm 0.2$ | 81$\pm 2$ | 5.52$\pm 0.11$ | 0.097 | 9.722 | 105.195 | 12.05 | 49 |
| | | 6 | 0.021<br>$\pm 0.003$ | 1.2$\pm 0.2$ | 77$\pm 2$ | 5.22$\pm 0.08$ | 0.160 | 13.570 | 49.146 | 12.87 | 65 |
| 2 | 9 | 3 | 0.0056<br>$\pm 0.0006$ | 2.7$\pm 0.4$ | 92$\pm 3$ | 6.1$\pm 0.2$ | 0.049 | 6.110 | 184.571 | 10.51 | 34 |
| | | 6 | 0.017<br>$\pm 0.004$ | 0.8$\pm 0.2$ | 84$\pm 2$ | 4.85$\pm 0.07$ | 0.266 | 22.669 | 59.364 | 11.83 | 103 |
| 3 | 4 | 3 | 0.0054<br>$\pm 0.0014$ | 7$\pm 2$ | 77$\pm 7$ | 6.3$\pm 0.5$ | 0.015 | 2.140 | 201.661 | 11.95 | 12 |
| | | 6 | 0.020<br>$\pm 0.003$ | 1.2$\pm 0.2$ | 85$\pm 4$ | 5.3$\pm 0.2$ | 0.112 | 15.136 | 50.156 | 11.58 | 74 |

**Figure 3.**
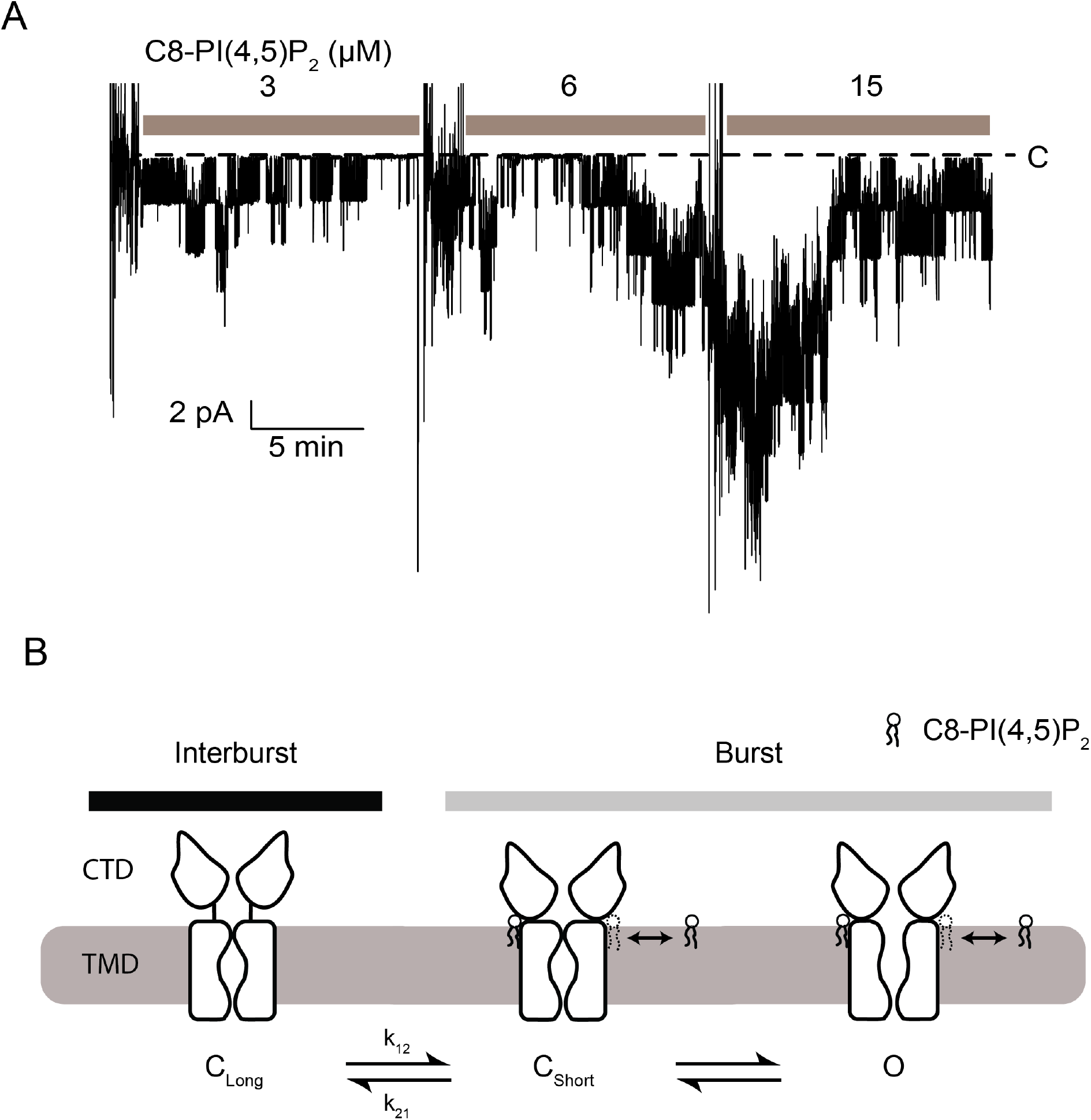
Single-channel analysis of Kir2.2 at various C8-PI(4,5)P_2_ concentrations. (A) Recording from a bilayer with multiple Kir2.2 channels in the presence of 3, 6 and 15 μM C8-PI(4,5)P_2_. Large current deflections between adjacent C8-PI(4,5)P_2_ concentration are due to mixing of the solution after each addition of C8-PI(4,5)P_2_. The membrane was held at −100 mV. Baseline is indicated as dashed line. (B) Schematic of the Kir2.2 gating model. The quiescent intervals correspond to the CTD-undocked, closed conformation. After binding enough C8-PI(4,5)P_2_, Kir2.2 will enter a burst, which corresponds to the CTD-docked conformation. The slow kinetic process of transition between the burst and quiescent states (k_12_ and k_21_) is sensitive to C8-PI(4,5)P_2_. Within the burst, the channel can rapidly transit between closed and open states. These rapid gating transitions are insensitive to C8-PI(4,5)P_2_ concentration. After a sufficient number of PI(4,5)P_2_ molecules have bound to enter a burst, at least one more can still bind. Therefore, lipids that bind to the channel can exchange within the burst.

The functional analysis leads to the simple conclusion that only the slow kinetic process of transition between the burst and quiescent states is sensitive to C8-PI(4,5)P_2_. The structural studies show that the binding of C8-PI(4,5)P_2_ is associated with a large conformational change between the CTD-undocked and CTD-docked structures (Hansen et al., 2011; Niu et al., 2020; Tao et al., 2009). We thus propose that the slow gating transitions in the channel recordings correspond to CTD engagement and disengagement, and that when the CTD is engaged, the pore can open (Figure 3B). The rapid gating transitions within a burst would then represent C8-PI(4,5)P_2_-independent conformational changes that occur elsewhere along the ion conduction pathway. According to this model, the channel record provides a dynamic read-out of the CTD engagement and disengagement process, whose equilibrium is shifted by the C8-PI(4,5)P_2_ concentration.

### Effect of phosphatidylinositol lipid derivatives on gating

Earlier studies have shown that phosphatidylinositol lipids with different phosphate substitutions can also interact with Kir channels, including Kir2.2 (Logothetis et al., 2015). Figure 4A shows a number of the chemical interactions between C8-PI(4,5)P_2_ and Kir2.2 derived from the crystal structure (Hansen et al., 2011). Individual membrane recordings using the planar bilayer system show the effects of several different phosphatidylinositol lipids (Figure 4 B-G). During each recording, application of the lipid under examination was followed by application of C8-PI(4,5)P_2_ to ensure the presence of Kir2.2 channels in the membrane. Soluble (C8) phosphatidylinositol lipids were used except for PI(5)P, which was only available in a long acyl-chain form and thus was included as part of the membrane’s lipid composition. We found that only phosphatidylinositol lipids with the 5’ phosphate can activate Kir2.2 channel and the magnitude of Kir2.2 activation for the lipid concentrations applied was PI(4,5)P_2_ ≈ PI(3,4,5)P_3_ > PI(3,5)P_2_ > PI(5)P (Figure 4 B-D). Essentially no activation was observed with PI(3)P, PI(4)P and PI(3,4)P_2_ (Figure 4 E-G). The results are consistent with a previous study showing that Kir2.2 can be activated in membrane patches from cells by C8-PI(4,5)P_2_ and C8-PI(3,4,5)P_3_, but not C8-PI(3,4)P_2_ (Rohacs et al., 2003). In the crystal structure of Kir2.2, the 5’ phosphate forms ionized hydrogen bonds with several basic residues at and near the base of the inner helix, which forms the gate (Figure 4A) (Hansen et al., 2011). This is compatible with the functional requirement of 5’ phosphate to open Kir2.2 channel.

**Figure 4.**
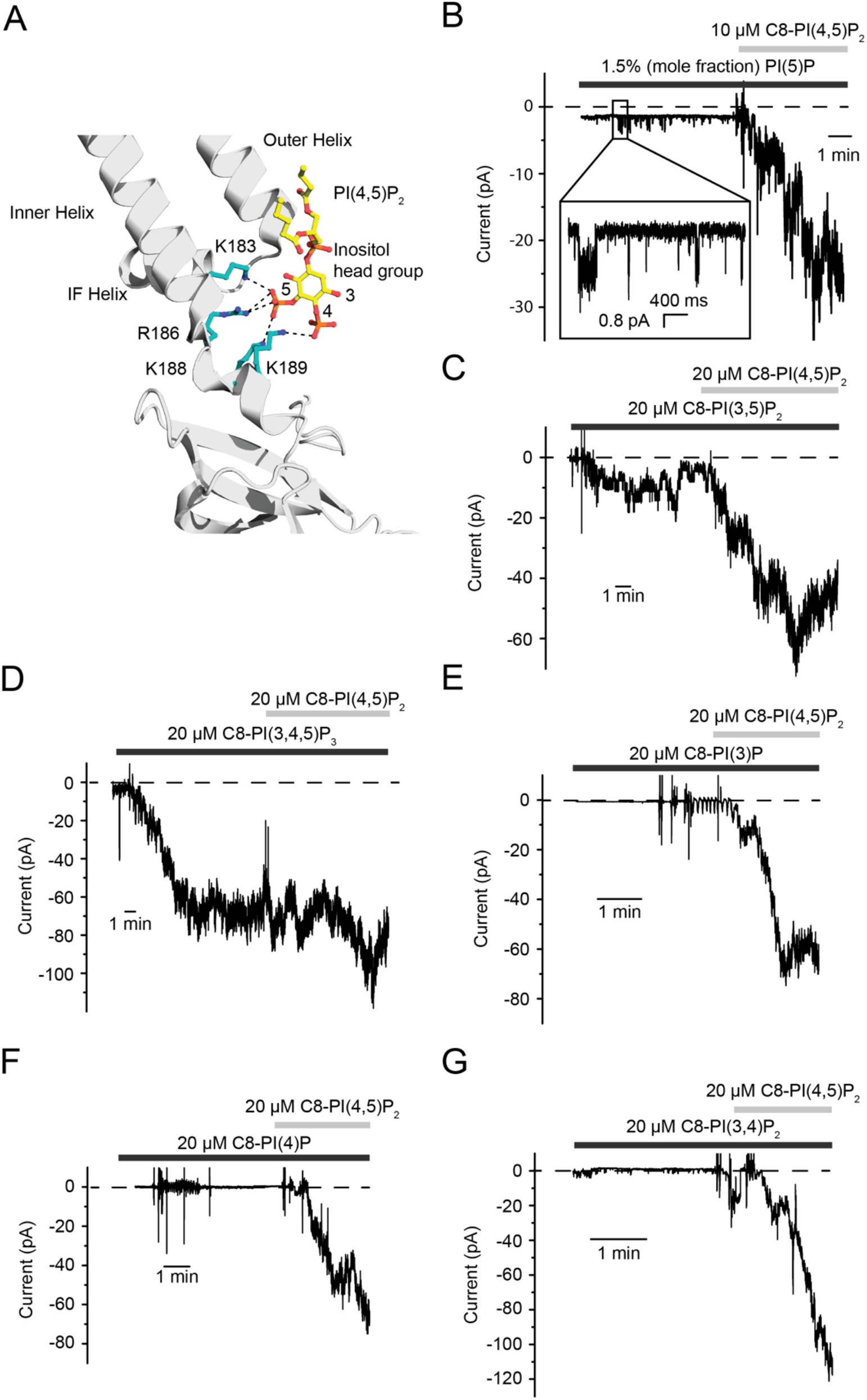
Effect of different phosphatidylinositol lipids on Kir2.2. (A) C8-PI(4,5)P_2_ binding site on Kir2.2 (PDB 3SPH). The channel is shown as grey ribbon. C8-PI(4,5)P2 is shown as sticks and colored according to atom type: oxygen, red; phosphorous, orange; and carbon, yellow. Sidechains of residues that form hydrogen bonds with 4’ or 5’ phosphate are shown as sticks and colored teal. (B) PI(5)P can activate Kir2.2. Short openings (inset) were observed from membranes with 1.5% (mole fraction) PI(5)P. Further application of 10 μM C8-PI(4,5)P_2_ resulted in a current increase. The membrane was held at −100 mV. (C) C8-PI(3,5)P2 can activate Kir2.2. Further application of 20 μM C8-PI(4,5)P_2_ resulted in a significant increase of current. The membrane was held at −100 mV. (D) C8-PI(3,4,5)P3 activates Kir2.2 to a similar extent as C8-PI(4,5)P_2_. The membranes were held at −100 mV. (E-G) 20 μM C8-PI(3)P (E), C8-PI(4)P (F) or C8-PI(3,4)P_2_ (G) failed to activate Kir2.2 despite the presence of Kir2.2 channels in the membrane, demonstrated by further addition of 20 μM C8-PI(4,5)P_2_. The membranes were held at −100 mV. Large current deflections are caused by mixing. Zero-current level is indicated as dashed line.

Do phosphatidylinositol lipids that do not activate Kir2.2 fail to bind altogether or do they bind but fail to open the gate? The data in Figure 5 suggest the latter, that certain phosphatidylinositol lipids inhibit Kir2.2 activation by competing with PI(4,5)P_2_ for its binding site. Following Kir2.2 activation by 6 μM C8-PI(4,5)P_2_, addition of 20 μM PI(3)P, PI(4)P or PI(3,4)P_2_ caused a pronounced reduction in current (Figure 5 A-C). PI(3,5)P_2_, which itself activates Kir2.2 but to a lesser extent than PI(4,5)P_2,_ also reduced current apparently by competition for the site (Figure 5D). In membranes with a small number of channels we tried to determine how a competing lipid, C8-PI(4)P, influences the kinetics of gating (Figure 5E and Table 2). Again, only the rate constants into and out of C_Long_ are affected, as if addition of C8-PI(4)P mimics a reduction in the concentration of C8-PI(4,5)P_2_.

**Table 2.** Kinetics of the C8-PI(4)P competition. Kinetics of Kir2.2 based on scheme 3 (C_Long_ == C_Short_ == O) in the presence of 6 μM C8-PI(4,5)P_2_ before and after application of C8-PI(4)P. Rate constant (k), open probability (P(o)), mean burst duration (T_burst_), mean inter-burst interval (T_inter-burst_) and mean number of short closures per burst (#C_Short_ per burst) are calculated from 3 different recordings.

| Recording Number | Channel Number | C8-PI(4,5)P<sub>2</sub>(μM) | C8-PI(4)P(μM) | $k_{12}$(s<sup>-1</sup>) | $k_{21}$(s<sup>-1</sup>) | $k_{23}$(s<sup>-1</sup>) | $k_{32}$(s<sup>-1</sup>) | P(o) | T<sub>burst</sub>(s) | T<sub>inter-burst</sub>(s) | Mean Duration of C<sub>Short</sub>(ms) | #C<sub>Short</sub>per burst |
| --- | --- | --- | --- | --- | --- | --- | --- | --- | --- | --- | --- | --- |
| 1 | 5 | 6 | - | 0.013<br>±0.002 | 0.90±0.14 | 52±2 | 4.20±0.11 | 0.134 | 14.907 | 77.772 | 19.04 | 57 |
| | | | 6 | 0.005<br>±0.002 | 13±4 | 57±8 | 3.0±0.5 | 0.006 | 1.865 | 266.014 | 14.36 | 4 |
| 2 | 4 | 6 | - | 0.007<br>±0.002 | 0.62±0.13 | 53±2 | 3.35±0.09 | 0.170 | 27.597 | 153.769 | 18.59 | 86 |
| | | | 6 | 0.0013<br>±0.0003 | 1.8±0.4 | 43±2 | 3.7±0.2 | 0.046 | 7.475 | 784.335 | 22.26 | 25 |
| 3 | 5 | 6 | - | 0.0027<br>±0.0007 | 0.14±0.05 | 43.9±1.2 | 3.72±0.08 | 0.235 | 94.889 | 370.114 | 22.72 | 324 |
| | | | 6 | 0.0005<br>±0.0005 | 0.39±0.09 | 42.1±1.2 | 3.03±0.08 | 0.117 | 38.427 | 2271.968 | 23.54 | 108 |

**Figure 5.**
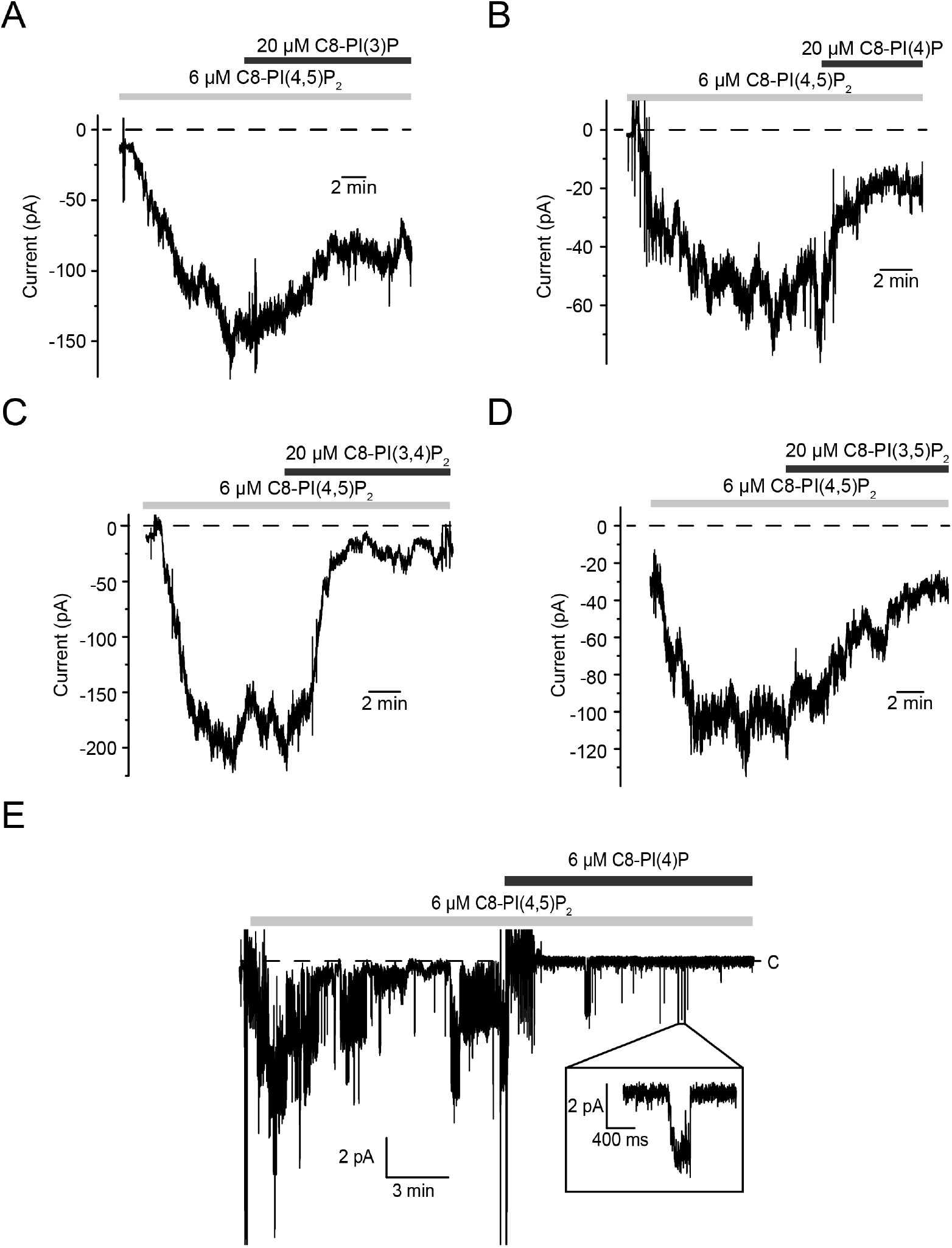
Competition by various phosphatidylinositol lipids with C8-PI(4,5)P_2_. (A-D) C8-PI(3)P (A), C8-PI(4)P (B), C8-PI(3,4)P_2_ (C) or C8-PI(3,5)P_2_ (D) inhibited Kir2.2 activation. Following Kir2.2 activation by 6 μM C8-PI(4,5)P_2_, application of 20 μM C8-PI(3)P (A), C8-PI(4)P (B), C8-PI(3,4)P_2_ (C) or C8-PI(3,5)P_2_ (D) caused a reduction in current. The membranes were held at −100 mV. (E) Current recorded from a bilayer containing multiple Kir2.2 channels after application of 6 μM C8-PI(4,5)P_2_ followed by additional 6 μM C8-PI(4)P. The membrane was held at −100 mV.

## Discussion

These results build on previous studies showing that PI(4,5)P_2_ is necessary and sufficient to open Kir2.2 (D’Avanzo et al., 2010). They also build on earlier characterizations of phosphatidylinositol lipid specificity in the activation of Kir channels in general (Logothetis et al., 2015). The present study advances our understanding by analyzing the phosphatidylinositol lipid-dependent gating of Kir2.2 in the compositionally-defined lipid bilayer system. It also presents a mechanistic model derived by correlating C8-PI(4,5)P_2_-dependent changes in the atomic structure of Kir2.2 with C8-PI(4,5)P_2_-dependent changes in the kinetics of gating. The accuracy of the kinetic data is limited for reasons described above. Nevertheless, the conclusion that C8-PI(4,5)P_2_ influences only the relatively slow transitions that govern exchange between the burst and quiescent periods is robust. In the model we connect the large conformational change observed in structural studies - C8-PI(4,5)P_2_-mediated engagement between the CTD and TMD – with the burst-quiescent period interconversions. This idea is depicted in cartoon form in Figure 3B. In the CTD-docked conformation, the pore opens and closes rapidly in a C8-PI(4,5)P_2_-independent manner. Other Kir channels also exhibit burst kinetics (Enkvetchakul et al., 2000; Jin et al., 2008) and in KATP, ATP-independent gating within burst periods has been attributed to processes inside the selectivity filter (Proks et al., 2001). The basis for rapid gating in Kir2.2 is still unknown.

A G protein-independent GIRK channel was found to exhibit a more complex dependence on PI(4,5)P_2_ than we describe here for Kir2.2 (Jin et al., 2008). In the mutant GIRK channel, in addition to influencing the burst (but not the inter-burst) duration, multiple open states were deduced and attributed to different degrees of PI(4,5)P_2_ occupation. In Kir2.2 we observe only a single open state and can only say the following regarding the functional stoichiometry of PI(4,5)P_2_ activation. That the quiescent periods shorten with increasing PI(4,5)P_2_ concentration indicates that at least one bound PI(4,5)P_2_ molecule is required to enter a burst. That the burst periods lengthen with increasing PI(4,5)P_2_ concentration indicates that after a sufficient number of PI(4,5)P_2_ molecules have bound to enter a burst, at least one more can still bind (Figure 3B). If this were not the case, then the burst duration would be independent of the PI(4,5)P_2_ concentration. In conclusion, somewhere between 1 and 3 PI(4,5)P_2_ molecules would seem to be required to stabilize the structure underlying the burst state, which we propose is the CTD-docked state. We know from the structures that 4 PI(4,5)P_2_ molecules can bind to the CTD-docked conformation, but if the model is correct, fewer than 4 can support the CTD-docked conformation.

The requirement of a 5’ phosphate to achieve channel opening seems compatible with the crystal structure of Kir2.2 with PI(4,5)P_2_ bound because the 5’ phosphate interacts directly with amino acids on the inner helix, which forms the gate. PI(4,5)P_2_ also makes other interactions with the channel so it is understandable that a phosphatidylinositol lipid without the 5’ phosphate would still bind to the site. This is undoubtedly why PI(4)P, for example, competes with PI(4,5)P_2_ for the phosphatidylinositol lipid binding site on the channel. PI(4)P as a competing lipid is particularly interesting because it, along with PI(4,5)P_2_, is abundant in the plasma membrane (Hammond and Balla, 2015). Moreover, PI(4)P is a precursor in the synthesis of PI(4,5)P_2_ and dephosphorylation of PI(4,5)P_2_ generates PI(4)P (Di Paolo and De Camilli, 2006; Hammond and Balla, 2015; Logothetis et al., 2015). Because PI(4,5)P_2_ activates and PI(4)P competitively inhibits, changes in lipid metabolism could give rise to a very steep change in the level of Kir2.2 activity.

## Materials and methods

### Key resources table

| Reagent type (species) or resource | Designation | Source or reference | Identifiers | Additional information |
| --- | --- | --- | --- | --- |
| Gene (Gallus gallus Kir2.2) | Kir2.2 | synthetic |  | Synthesized at Bio Basic, Inc. |
| Strain, strain background (Escherichia coli) | DH10Bac | ThermoFisher | 10361012 |  |
| Recombinant DNA reagent | cKir2.2-GFP-1D4 pEG BacMam | https://doi.org/10.7554/eLife.15750.001 |  | Maintained at the Mackinnon lab |
| Cell line (Spodoptera frugiperda) | Sf9 | ATCC | Cat# CRL-1711 |  |
| Cell line (Homo sapiens) | HEK293S GnTI<sup>-</sup> | ATCC | Cat# CRL-3022 |  |
| Chemical compound, drug | SF-900 II SFM medium | GIBCO | Cat# 10902–088 |  |
| Chemical compound, drug | L-Glutamine (100x) | GIBCO | Cat# 25030–081 |  |
| Chemical compound, drug | Pen Strep | GIBCO | Cat# 15140–122 |  |
| Chemical compound, drug | Grace's insect medium | GIBCO | Cat# 11605–094 |  |
| Chemical compound, drug | Freestyle 293 medium | GIBCO | Cat# 12338–018 |  |
| Chemical compound, drug | Fetal bovine serum | GIBCO | Cat# 16000–044 |  |
| Chemical compound, drug | Cellfectin II reagent | Invitrogen | Cat# 10362100 |  |
| Chemical compound, drug | n-Decyl-b-D-Maltopyranoside (DM) | Anatrace | D322S |  |
| Chemical compound, drug | 1,2-dioctanoyl-sn-glycero-3-phospho-(1'-myo-inositol-4',5'-bisphosphate) (ammonium salt) (C8-PI(4,5)P2) | Avanti Polar Lipids | 850185P |  |
| Chemical compound, drug | 1-stearoyl-2-arachidonoyl-sn-glycero-3-phospho-(1'-myo-inositol-5'-phosphate)<br>(ammonium salt)<br>(PI(5)P) | Avanti Polar Lipids | 850190P |  |
| Chemical compound, drug | 1,2-dioctanoyl-sn-glycero-3-phospho-(1'-myo-inositol-3',5'-bisphosphate)<br>(ammonium salt)<br>(C8-PI(3,5)P2) | Avanti Polar Lipids | 850184P |  |
| Chemical compound, drug | 1,2-dioctanoyl-sn-glycero-3-(phosphoinositol-3-phosphate)<br>(ammonium salt)<br>(C8-PI(3)P) | Avanti Polar Lipids | 850187P |  |
| Chemical compound, drug | 1,2-dioctanoyl-sn-glycero-3-phospho-(1'-myo-inositol-4'-phosphate)<br>(ammonium salt)<br>(C8-PI(4)P) | Avanti Polar Lipids | 850182P |  |
| Chemical compound, drug | 1,2-dioctanoyl-sn-glycero-3-phospho-(1'-myo-inositol-3',4'-bisphosphate)<br>(ammonium salt)<br>(C8-PI(3,4)P2) | Avanti Polar Lipids | 850183P |  |
| Chemical compound, drug | 1,2-dioctanoyl-sn-glycero-3-phospho-(1'-myo-inositol-3',4',5'-trisphosphate)<br>(ammonium salt) | Avanti Polar Lipids | 850186P |  |
| Chemical compound, drug | 1,2-dioleoyl-sn-glycero-3-phosphoethanolamine<br>(DOPE) | Avanti Polar Lipids | 850725P |  |
| Chemical compound, drug | 1-palmitoyl-2-oleoyl-glycero-3-phosphocholine<br>(POPC) | Avanti Polar Lipids | 850457C |  |
| Chemical compound, drug | 1-palmitoyl-2-oleoyl-sn-glycero-3-phospho-L-serine<br>(sodium salt) (POPS) | Avanti Polar Lipids | 840034P |  |
| Chemical compound, drug | 1-palmitoyl-2-oleoyl-sn-glycero-3-phosphoethanolamine (POPE) | Avanti Polar Lipids | 850757P |  |
| Chemical compound, drug | 1-palmitoyl-2-oleoyl-sn-glycero-3-phospho-(1'-rac-glycerol) (sodium salt) (POPG) | Avanti Polar Lipids | 840457C |  |
| Commercial assay or kit | Superdex 200 Increase 10/300 GL | GE Healthcare Life Sciences | 28990944 |  |
| Software, algorithm | Fit | doi: [10.1016/S0006-3495(00)76636-7](https://doi.org/10.1016/S0006-3495%2800%2976636-7) |  |  |
| Software, algorithm | pClamp | Axon Instruments, Inc |  |  |
| Software, algorithm | Mathematica | Wolfram | SCR_014448 |  |
| Software, algorithm | Pymol | PyMOL Molecular Graphics System, Schrödinger, LLC | http://www.pymol.org |  |
| Software, algorithm | Clampfit | Molecular Devices |  |  |

### Cloning, expression, and purification

A synthetic gene fragment (Bio Basic, Inc.) encoding residues 38 to 369 of chicken Kir2.2 (cKir2.2) channel (GI:118097849) was subcloned into a modified pEG BacMam vector with a C-terminal green fluorescent protein (GFP)-1D4 tag linked by a preScission protease site (Goehring et al., 2014). This construct was used in all experiments in this study.

Bacmid containing cKir2.2 gene was generated according to the manufacturer’s instructions (Invitrogen) by transforming the cKir2.2 pEG BacMam construct into E. coli DH10Bac cells. The bacmid was then transfected into *Spodoptera frugiperda* Sf9 cells to produce baculoviruses using Cellfectin II (Invitrogen). After two rounds of amplification, P3 viruses were added to HEK293S GnTI^−^ cells (ATCC) in a 1:10 (v:v) ratio for protein expression. Suspension cultures of HEK293S GnTI^−^ cells were grown in Freestyle 293 media (GIBCO) supplemented with 2% FBS (GIBCO) at 37°C to a density around 1.5-3×10^6^ cells/ml. After incubating the infected cell for 20 hours at 37°C, 10 mM sodium butyrate was added and the temperature was changed to 30°C. Cells were harvested ~40 hours after changing the temperature (Goehring et al., 2014). 4L cell pellet was first resuspended in 200 mL hypotonic lysis buffer (20 mM Tris-HCl pH 8, 1 mM EDTA, 1 mM phenylmethysulfonyl fluoride (PMSF), 0.1 mg/ml 4-(2-Aminoethyl) benzenesulfonyl fluoride hydrochloride (AEBSF), 0.1 mg/ml soybean trypsin inhibitor, 1 mM benzamidine, 1 μg/ml pepstatin A, 1 μg/ml leupeptin, 5 μg/ml aprotinin and 0.02 mg/ml Dnase I) at 4°C. The lysate was then centrifuged and the pellet was homogenized using douncer into 100 mL extraction buffer containing 20 mM Tris-HCl pH 8, 1 mM EDTA, 320 mM KCl, 2 mM PMSF, 0.1 mg/ml AEBSF, 0.1 mg/ml soybean trypsin inhibitor, 1 mM benzamidine, 1 μg/ml pepstatin A, 1 μg/ml leupeptin, 5 μg/ml aprotinin and 0.1 mg/ml Dnase I. The lysate was supplemented with 40 mM DM to extract at 4°C for 1-2 hrs and then centrifuged. The supernatant was incubated with GFP nanobody-conjugated affinity resin (CNBr-activated Sepharose 4B resin from GE Healthcare) by rotating at 4°C for 1 hour. Resin was then washed with 10 column volumes of wash buffer (20 mM Tris-HCl pH 8, 1 mM EDTA, 150 mM KCl and 6 mM DM). PreScission protease (~1:20 w:w ratio) was then added for digestion overnight by rotating. Flow through was collected, concentrated and loaded onto Superdex 200 increase size exclusion column (GE Healthcare) equilibrated in 20 mM Tris-HCl pH 8, 1 mM EDTA, 150 mM KCl, 4 mM DM, 10 mM DTT and 2 mM TCEP. The purified protein was then concentrated for reconstitution.

### Reconstitution of proteoliposomes

The reconstitution was performed as previously described (Brohawn et al., 2012; Heginbotham et al., 1999; Long et al., 2007; Tao and MacKinnon, 2008; Wang et al., 2014) with minor modifications. Briefly, a lipid mixture composed of 3:1 (w:w) 1-palmitoyl-2-oleoyl-sn-glycero-3-phosphoethanolamine (POPE) : 1-palmitoyl-2-oleoyl-sn-glycero-3-phospho-(1′-rac-glycerol) (POPG) (Avanti) was dried under Argon, rehydrated in reconstitution buffer (10 mM potassium phosphate pH 7.4, 150 mM KCl, 1 mM EDTA and 3 mM DTT) to 20 mg/ml by rotating for 20 min at room temperature followed by sonication with a bath sonicator. 1% DM was then added. The lipid mixture was rotated for 30 min and sonicated again till clear. Equal volume of protein (at 2 mg/ml and 0.2 mg/ml) and lipid (at 20 mg/ml) were mixed, resulting in protein: lipid (w:w) ratios of 1:10 and 1:100 respectively. The mixture was incubated at 4°C for 1 hour and then dialyzed against 2L reconstitution buffer for 2 days, exchanging buffer every 12 hours. Biobeads was added to the reconstitution buffer for the last 12 hours. The resulting proteoliposomes were frozen with liquid nitrogen and stored at −80°C.

### Electrophysiology

The bilayer experiments were performed as previously described with minor modifications (Ruta et al., 2003; Wang et al., 2014). A piece of polyethylene terephthalate transparency film separates the two chambers of a polyoxymethylene block which are filled with buffer containing 10 mM potassium phosphate pH 7.4, 150 mM KCl and 2 mM Mg^2+^.

A 20 mg/ml decane lipid mixture (w:w:w 2:1:1) of 1,2-dioleoyl-sn-glycero-3-phosphoethanolamine (DOPE) : 1-palmitoyl-2-oleoyl-sn-glycero-3-phosphocholine (POPC) : 1-palmitoyl-2-oleoyl-sn-glycero-3-phospho-L-serine (POPS) (Avanti) was pre-painted over a ~100 μm hole on the transparency film. Voltage was controlled with an Axopatch 200B amplifier in whole-cell mode. The analog current signal was low-pass filtered at 1 kHz (Bessel) and digitized at 10 kHz with a Digidata 1322A digitizer. Digitized data were recorded on a computer using the software pClamp (Molecular Devices, Sunnyvale, CA). Experiments were performed at room temperature. For macroscopic current recordings, data reduction with a reduction factor of 100 and 5 Hz Gaussian low-pass filter was applied for plotting purpose. For single-channel recordings, a 300 Hz Gaussian low-pass filter was applied to the expanded trace (Figure 2A, inset), or data reduction with a reduction factor of 200 was applied to the whole single-channel recording for plotting.

### Kinetic analysis

Recordings containing 1-9 channels were idealized through half-amplitude threshold crossing and analyzed using Clampfit software (Molecular Devices).

For single-channel recordings, open or closed dwell time distributions for events within the burst were fitted to an exponential probability density function. Models with different term numbers were compared and the best model was selected (Horn, 1987).

Multi-channel recordings were analyzed as described (Csanady et al., 2000). Eventlists were fitted using a three-state linear 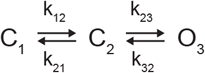 scheme. Dead time was 0.27 ms. Channel numbers used in the multi-channel kinetic analysis were estimated using the maximum number of observed open channel levels (from 15 μM C8-PI(4,5)P_2_ recordings). Rate constants between all states (k_12_, k_21_, k_23_ and k_32_) were obtained by simultaneous fit to the dwell-time histograms of all conductance levels (Csanady, 2000). The mean number of short closures per burst, mean burst and inter-burst duration were then calculated based on the rate constants -- mean number of short closures per burst = 1 + k_23_ / k_21_, mean inter-burst duration = 1 / k_12_, and mean burst duration = (1 / k_21_) (1+ k_23_ / k_32_).

### Estimation of the channel number used in multi-channel kinetic analysis

To evaluate the validity of using the maximum number of observed open channel levels (from 15 μM C8-PI(4,5)P_2_ recordings) as the channel number (N) in the multi-channel kinetic analysis, we ask what is the probability of observing N channels at least once during the duration of the record? This probability is a function of N, the rate constants, and the initial condition. This probability is given by the integral of the first passage time probability density function for the appearance of the right-most state for the scheme, for N = 8, shown here:

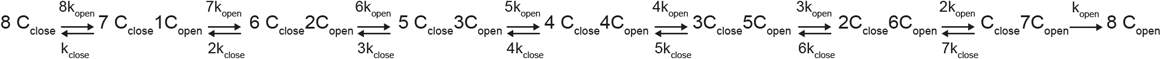

This scheme, with the rate constants weighted as shown, models a membrane with N = 8 identical channels. The quiescent state of each channel is denoted as C_close_ and the burst is denoted as C_open_. The left-most state represents the membrane with 8 closed channels. The right-most state represents the membrane with 8 open (i.e. burst channels).

Using Mathematica (Wolfram), we estimate that the mean first passage time is 268 s and that the probability of observing 8 channels at least once during the duration of the record (10 min) is 0.9. Given the 10% chance that we cannot see all 8 channels open within 10 min an the problem of channel disappearance over time, it is possible that we have underestimated the channel number. We therefore ask, if we assign the incorrect value for N, will our conclusion that C8-PI(4,5)P_2_ affects only the burst and inter-burst periods be wrong? Using recording 3 (Table 1), in which we have assigned N = 4 on the basis of direct observation we re-analyzed the record for N = 4 to 10 (Table 3). The value of N had little influence on the determination of kinetic values within the burst and had a small influence on k_21_ and mean burst duration. The main affect was on k_12_ and mean inter-burst duration. But importantly, when C8-PI(4,5)P_2_ is increased, the burst periods lengthen and the quiescent periods shorten. Thus, our general conclusion that C8-PI(4,5)P_2_ concentrations influence the transitions between the burst and inter-burst states holds even if we have underestimated N.

**Table 3.** Kinetics of Kir2.2 with different channel numbers. Re-analysis of Recording 3 from Table 1, using various channel numbers (N).

| Recording Number | Channels Number | C8-PI(4,5)P<sub>2</sub>(μM) | k<sub>12</sub>(s<sup>-1</sup>) | k<sub>21</sub>(s<sup>-1</sup>) | k<sub>23</sub>(s<sup>-1</sup>) | k<sub>32</sub>(s<sup>-1</sup>) | P(o) | T<sub>burst</sub>(s) | T<sub>inter-burst</sub>(s) | Mean Duration of C<sub>Short</sub>(ms) | #C<sub>Short</sub> per burst |
| --- | --- | --- | --- | --- | --- | --- | --- | --- | --- | --- | --- |
| 3 | 4 | 3 | 0.0054<br>±0.0014 | 7±2 | 77±7 | 6.3±0.5 | 0.015 | 2.140 | 201.661 | 11.95 | 11.52 |
|  |  | 6 | 0.020<br>±0.003 | 1.2±0.2 | 85±4 | 5.3±0.2 | 0.112 | 15.136 | 50.156 | 11.58 | 73.92 |
|  | 5 | 3 | 0.0044<br>±0.0016 | 7±2 | 75.8±7.1 | 6.2±0.6 | 0.0117 | 2.110 | 246.782 | 12.12 | 11.30 |
|  |  | 6 | 0.016<br>±0.003 | 1.3±0.3 | 86±4 | 5.33±0.19 | 0.0892 | 13.353 | 63.729 | 11.43 | 66.13 |
|  | 6 | 3 | 0.0036<br>±0.0014 | 7±2 | 76±7 | 6.2±0.6 | 0.0098 | 2.096 | 300.129 | 12.12 | 11.22 |
|  |  | 6 | 0.013<br>±0.003 | 1.4±0.3 | 87±4 | 5.4±0.2 | 0.0743 | 12.551 | 78.305 | 11.37 | 62.42 |
|  | 7 | 3 | 0.0031<br>±0.0009 | 7±2 | 76±7 | 6.2±0.5 | 0.0084 | 2.085 | 353.833 | 12.12 | 11.15 |
|  |  | 6 | 0.011<br>±0.002 | 1.4±0.3 | 87±4 | 5.37±0.17 | 0.0637 | 12.036 | 92.779 | 11.33 | 60.01 |
|  | 8 | 3 | 0.0027<br>±0.0013 | 7±2 | 76±7 | 6.2±0.5 | 0.0073 | 2.077 | 407.228 | 12.12 | 11.11 |
|  |  | 6 | 0.0095<br>±0.0015 | 1.5±0.3 | 87±4 | 5.39±0.18 | 0.0557 | 11.678 | 107.161 | 11.31 | 58.34 |
|  | 9 | 3 | 0.0024<br>±0.0006 | 7±2 | 76±7 | 6.2±0.6 | 0.0065 | 2.072 | 460.095 | 12.12 | 11.08 |
|  |  | 6 | 0.0084<br>±0.0016 | 1.5±0.3 | 87±4 | 5.4±0.2 | 0.0496 | 11.438 | 121.653 | 11.27 | 57.25 |
|  | 10 | 3 | 0.0021<br>±0.0009 | 7±2 | 76±7 | 6.2±0.6 | 0.0059 | 2.067 | 513.606 | 12.12 | 11.05 |
|  |  | 6 | 0.0075<br>±0.0015 | 1.5±0.3 | 87±4 | 5.4±0.2 | 0.0446 | 11.247 | 136.103 | 11.26 | 56.36 |

## Acknowledgements

We thank Laszlo Csanady for sharing his software for the multi-channel kinetic analysis. We thank members of the MacKinnon laboratory, especially Chen Zhao and Jesper Levring (Chen Lab, Rockefeller University) for helpful discussions. Thank Chen Zhao, Jesper Levring, Maria Falzone, James Lee (Chen Lab, Rockefeller University) and Venkata Shiva Mandala for advice on the manuscript.

This work was supported in part by GM43949. RM is an investigator in the Howard Hughes Medical Institute.

---

**Figure 1-Figure Supplement 1.**
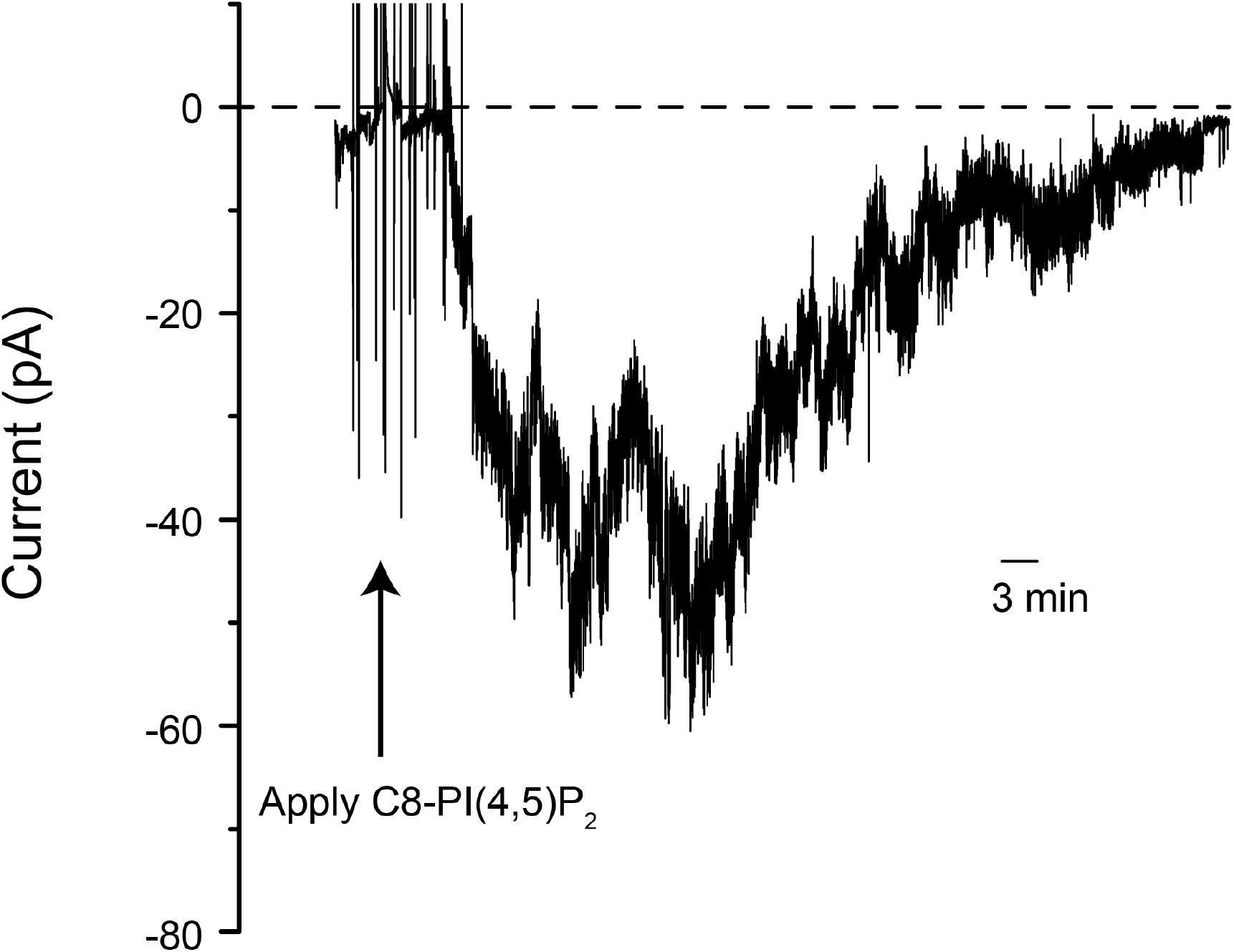
Disappearance of channel activity. C8-PI(4,5)P_2_-induced current begins to decrease spontaneously after about 30 min. Representative current trace before and after addition of 4 μM C8-PI(4,5)P_2_ to the top chamber is shown. Zero-current level is indicated by dashed line.

